# ^1^Cross Sectional Study of Middle East Respiratory Syndrome (Mers-Cov Infection) in Camels at Selected Sites of Amibara District, Afar Region, Ethiopia

**DOI:** 10.1101/2020.10.25.353227

**Authors:** Demeke Sibhatu, Gezahagne Mamo, Fasil Aklillu, Demeke Zewde, Elias Walelign, Ayelech Muluneh, Abdi Aliey, Tadele Mirkena, Nega Tewolde, Getachew Gari, Gijs van ‘t Klooster, Ihab Elmasry, Sophie VonDobschuetz, Malik Peiris, Daniel Chu, Ranawaka APM Perera, Yilma Jobire

## Abstract

**Background:** A Cross sectional study of Middle East Respiratory Syndrome Corona virus (MERS-CoV) in Camel was conducted between February 2018 to April 2019 in three selected sites of Amibara district of Afar region, Northeast Ethiopia. The study was aimed to observe the current sero-prevalence status of MERS-CoV, assess the presence of active cases through detection RNA Viral particle and investigate possible risk factors of MERS-CoV in camels. A total of 589 sera were collected and tested with indirect Enzyme linked ImmunoSorbent Assay (iELISA).

**Result:** The overall seroprevalance of MERS-CoV was 87.3% (n=514/589, 95% CI: 84.5-89.9). Association of different risk factors with seroprevalance revealed that origin (X^2^=13.39,P=0.001), sex (X^2^=4.5 P=0.034), age ((X^2^=185.7, P=0.001) season (X^2^=41.7, P=0.000) and reproduction status (X^2^=96.1, P=0.001) displayed a statistical significant difference among the groups (P<0.05) while herd size did not show a Significant difference among groups (p>0.05). In multivariable logistic regression analysis, age (OR=7.39, 95% CI:3.43-15.91), season (OR=4.83, 95% CI:-2.14-10.90), and in adult female camel reproduction status (OR=7.39,95% C I:3.43-15.91) showed statistically significant difference among the groups for MERS CoV antibody detection while risk factors of origin, animal sex and herd size difference were statistically insignificant. A total of 857 nasal swab samples were collected for the detection of MERS-CoV RNA particle. However, all swab samples tested by Real-time reverse transcription polymerase chain reaction (RT-PCR) technique were Negative for the virus.

**Conclusion:** In conclusion, the present study revealed a high seroprevalance of MERS CoV in adult camels. However, in spite of high seroprevalance the lack of any RNA viral particle in the study suggests the need for further in depth longitudinal study to detect the circulating virus focusing on juveniles and young camels whereby seroprevalance of antibody is low when compared with adult camel in order to get the active virus before the camel develop antibody. Moreover, the zoonotic significance and potential transmission routes of MERS CoV to pastoral communities should also be investigated and design strategy for the preparedness in control of the diseases in Ethiopia.

## INTRODUCTION

The one-humped camel (*Camelus dromedaries*) is an important livestock species exceptionally adapted to hot, dry and harsh environment due to heat and water deprivation tolerance. These tolerances in camels appear to be due to behavioral response that reduces heat absorption, a relatively efficient sweating mechanism for heat dissipation, an ability to reduce fecal and urine water loss and the ability to vary body temperature substantially. It is used for milk and meat production, transportation, and draught power **[1].** Camels are widely distributed in Ethiopian lowlands especially in Afar, Somali and Oromia region where by pastoralism is the dominant mode of life and mobility is an inherent strategy to efficiently utilize the spatially and temporally distributed pasture and water resources. Usually, large numbers of camels and other domestic animals from many different herds/flocks congregate at watering sites, and this may create a perfect condition for disease transmission and spread among animals. The same water sources are also shared by multitudes of wild animals **[2]**. According to CSA 2016/17 report, the camel population of dromedaries in Ethiopia is estimated to be about 1,209,321. Afar region has 474,146 camels **[3].**

Middle East Respiratory Syndrome (MERS) is a viral respiratory diseases within the largest group of Corona viruses (CoVs) belonging to Nidovirale order which includes Coronaviridae, Arteriviridae and Ro**n**aviridae families. The coronavirinae are further divide into four groups the alpha, beta, gamma and delta coronaviruses. MERS CoV is within beta corona virus group **[4]**. Dromedary camels are sturdily suspected of acting as a zoonotic source for human cases of MERS-CoV, by either direct contact through droplet infection via mucous membranes or indirect contact through milk, meat or urine. According to, Miguel *et al.*, (2016) five major points reason out accounts that suggest dromedary camels can play an important role in the epidemiology of MERS-CoV, possibly as a reservoir host:

- Corona-viruses are widespread in the animal kingdom (in bats and livestock), but MERS-CoV does not infect many of the hosts (e.g. sheep, goats, cattle, chickens, water buffaloes, birds, horses and) whereas high levels of sero positivity have been observed in dromedary camelids, ranging from 0% in Asia to as much as 100% in Africa and the Arabian Peninsula (with mean of 79%);
- The Mers-Cov isolated from dromedaries are genetically and phenotypically very similar to those infecting humans;
- Retrospective serological studies in Africa going back more than 30 years indicate long-term circulation of the virus in dromedary camels;
- Infection in dromedaries causes no or only mild respiratory symptoms, making it difficult to detect;
- Mers-Cov genome has likely undergone numerous recent recombination’, which suggests frequent co-infection, probably in camels, with distinct lineages of Mers-Cov **[5]**.

Studies have demonstrated that dromedary camels can act as a source of human MERS-CoV infection. Indeed, the current state of knowledge indicates that dromedary camels are the only animal species for which there is convincing evidence that they act as host species for Mers-Cov and hence a potential source of human infections **[6]**. Nonetheless, the route of infection of MERS CoV and types of exposures remain largely unknown, and only a small proportion of the primary cases have reported contact with camels. Other possible sources and vehicles of infection include food-borne transmission such as unpasteurized camel milk and raw meat, and medicinal use of camel urine **[7]**. Clearly, transmission from camels to humans does take place, and camel exposure is a risk factor for human infection, but such transmission is not efficient and infection is not directly proportional to exposure while in the other hand, many patients with clinically diagnosed MERS did not have an obvious history of direct exposure to camels or their products **[8]**.

Researchers found high percentages of animals sampled from Nigeria and Ethiopia being seropositive for Mers-Cov with an overall seropositivity of 94% in adult dromedaries in Nigeria and 93% and 97% for juvenile and adult animals, respectively, in Ethiopia **[9]**. More recently, **[10]** other researchers displayed a high seropositivity of 99.4% in camel of Ethiopia and also relatively higher Mers-Cov RNA detection in Ethiopia (15.7%) than in Burkina Faso (12.2%) and Morocco (7.6%). Also 10.6% virus detection rate observed by a study in Ethiopia as described by journals **[11].**

Other authors also described 93% seropositivity and 7% (n =7/100) MERS CoV RNA detection in Ethiopia, Afar region camels **[12]**. However, data from experimental camel infections conducted in the Middle East suggest that Mers-Cov causes only mild respiratory infection in camels **[13]**. Also study in Ethiopia between 2010-2011 reported 93-97% seropositivity **[9]**.

In Ethiopia, in spite of the high prevalence of Mers-Cov antibodies in camel as indicated in different studies, no human case has been reported to date, and only few ongoing studies have been carried out to investigate public health significance of MERS in highly exposed pastoralist community of Ethiopia who have close contact with camels requires serious attentions for further surveillance both for camel and exposed human population. So based on the mentioned points the objectives of the study were:

- To determine the current seroprevalance of MERS-CoV in camels with in selected sites of Amibara district, Afar Region.
- To identify the potential risk factors for MERS CoV in camels in order to control the disease.
- To detect and characterize MERS CoV from nasal swab of camels in the study sites.

## RESULT

### Sero-prevalence of MERS CoV antibody

Based on Indirect ELISA test results the overall prevalence of MERS CoV antibody in camels at study sites was 87.3% (n=514/589) (95% CI:84.5-89.9%). Association of different risk factors to seropositivity status of camels using X^2^ analysis revealed that there was a statstically significant difference in proportion of MERS Cov antibody positivity among the three study sites (X2 =13.7, p=0.001); Age categories (X2 = 185.69, p=0.000); sex categories (X2 = 4.5,p=0.034) and season (X2 = 41.69, p=0.000); and in reproduction status of adult female (X2 = 96.13, p=0.000); while no statistical significant difference were observed between herd sizes (X2 = 5.88, p=0.053) as illustrated in table 1.\

**1:** Association of different risk factors to seropositivity of camels MERS-CoV

| **Risk factors** | **No of tested camel** | **No of Positive** | **Prevalence (%positive)** | **X<sup>2</sup> value** | **p-value** |
| --- | --- | --- | --- | --- | --- |
| **Origin** |  |  |  | 13.39 | 0.001 |
| Andido | 289 | 266 | 92 |  |  |
| Melka sedi | 149 | 127 | 85.2 |  |  |
| Angelele | 151 | 121 | 80.1 |  |  |
| **Sex** |  |  |  | 4.50 | 0.034 |
| Male | 55 | 43 | 78.2 |  |  |
| Female | 534 | 471 | 88.2 |  |  |
| **Herd size** |  |  |  | 5.88 | 0.053 |
| Small | 100 | 83 | 83. |  |  |
| Medium | 168 | 155 | 92.3 |  |  |
| Large | 321 | 276 | 86 |  |  |
| **Age** |  |  |  | 185.69 | 0.001 |
| Juvenile | 89 | 39 | 43.8 |  |  |
| Young | 123 | 108 | 87.8 |  |  |
| Adult | 377 | 367 | 97.3 |  |  |
| **Season** |  |  |  | 41.69 | 0.001 |
| **Winter**<br>(December –February) | 88 | 79 | 89.8 |  |  |
| **Autumn** (September-November) | 272 | 260 | 95.6 |  |  |
| **Summer**<br>(June –August) | 229 | 175 | 76.4 |  |  |
| Dry | 162 | 157 | 96.9 |  |  |
| Pregnant | 68 | 64 | 94.1 |  |  |
| Lactating | 146 | 145 | 99.3 |  |  |

In multivariable logistic regression analysis young age (OR=7.39, 95% CI: 3.43-15.91), season from September-November (OR=4.83, 95% CI: 2.145-10.90), and in adult female camel lactation status ((OR=10.75, 95% CI: 1.15-100.08)) showed a statistically significant difference among the groups for MERS CoV antibody detection while risk factors of origin, animal sex and herd size did not show a statistical significant difference as indicated in table 2.

**Table 2:** Multivariable Logistic regression analysis of MERS CoV prevalence

| Risk factor | No of tested | No of positive and prevalence (%) | Crude OR (95% CI) | Adjusted OR (95% CI) |
| --- | --- | --- | --- | --- |
| **Origin** |  |  |  |  |
| Angelele | 151 | 121 (80.1) | 1 | 1 |
| Melka sedi | 149 | 127( 85.2) | 1.43(0.78-2.62) | 1.03(0.44-2.41) |
| Andido | 289 | 266 (92) | 2.87(1.60-5.14) | 1.00(0.42-2.41) |
| **Sex** |  |  |  |  |
| Male | 55 | 43 (78.2) | 1 | 1 |
| Female | 534 | 471 (88.2) | 2.09(1.04-4.16) | 0.86(0.35-2.14) |
| **Herd size** |  |  |  |  |
| Small | 100 | 83 (83) | 1 | 1 |
| Medium | 168 | 155( 92.3) | 2.44(1.13-5.27) | 2.35(0.89-5.15) |
| Large | 321 | 276 (86) | 1.26(0.68-2.31) | 1.51(0.71-3.55) |
| **Age** |  |  |  |  |
| Juvenile | 89 | 39 (43.8) | 1 | 1 |
| Young | 123 | 108 (87.8) | 9.23(4.55-18.28) | 7.39(3.43-15.91)* |
| Adult | 377 | 367 (97.3) | 47.05(22.11-100.10) | 21.91(0.27-1743.85) |
| **Season** |  |  |  |  |
| **Summer** ( June – August) | 229 | 175 (76.4) | 1 | 1 |
| **Autumn** ( September –November) | 272 | 260 (95.6) | 5.59(3.47-12.85) | 4.83(2.14-10.90)* |
| **Winter** (December – February) | 88 | 79 (89.7) | 2.71(1.27-5.76) | 4.10(1.30-12.86)* |
| **Production Status(Females)** |  |  |  |  |
| Pregnant | 68 | 64 (94.1) | 1 | 1 |
| Lactating | 146 | 145 (99.3) | 9.06(0.99-82.59) | 10.75(1.15-100.08) |
| Dry | 162 | 157 (96.9) | 1.95(0.51-7.54) | 2.49(0.53-9.92) |

\*Note: Risk factors displaying significant difference in Multivariable Logistic regression

### Viral RNA detection

All tested nasal swabs samples were negative for MERS CoV RNA particle by Real time polymerase chain reaction (RT-PCR) both at NAHDIC, Ethiopia and Hong Kong University (HKU).

## DISCUSSIONS

Middle East respiratory syndrome (MERS) is a zoonotic disease of global health concern, and dromedary camels are the source of human infection. In Ethiopia, a high seroprevalance of MERS-CoV in camel have been reported ranging from 93-97% in pastoral camel rearing areas of the country **[9]**. In the current study, a high prevalence of MERS-CoV with 87.3% (n=514/589), *(95% CI: 84.5-89.9)* was observed in camels of Amibara district, Afar Region. This high seroprevalance result was in agreement with previous studies in pastoral areas of Ethiopia who reported 85.1-99.4% in camels of Afar and Oromia **[5]**, 92.3% in Afar [12] 93-97% in Afar, Somali and Oromia regions **[9]**.

In multivariate logistic regression analysis three significant factors were observed in MERS CoV prevalence. Age; OR=7.39 (95%CI 3.43-15.91) with in this factor Adults >3 year are with high prevalence 97.3 %, young camels 1-3 years 87.8 % and Juvenile <1 year age 43.8 %. This study agree with previous study done by **[10]** in which antibodies detection rates were higher in older animals while Viral RNA was higher in young camels whereby they are free from antibody.

The reproduction status of female camels showed a considerable variation with OR=10.75(95%CI 1.15-100.08).With this result pregnant camels were being sated with low sero prevalence 94.1% when comparing with, Dry (96.9%) and lactating camels with (99.3 %). From this analysis we observe that high seroprevalance antibodies prevail in lactating camels when comparing with pregnant camels **[10]**.

Seasonal variation observed in this study (OR =4.83) illustrate high sero-prevalence is prevailed (95.6%) in autumn (September, October and November); (76.4%) in summer (June, July and August) and 89.7% winter (December, January and February). The high prevalence in autumn was due to gathering of camels at one place for prolonged period for the reason that camels are getting sufficient vegetation and grass. For this reason there had been high probability of infection and which induces he development of natural infection antibody. In winter the prevalence is low due to camels are dispersing far places in search of feed and water due to scarce of feed at one place. In this season the possibility of close contact and getting the disease through aerosol and developing antibody is limited.

Regarding seasonal factors, high seroprevalance was recorded in Autumn (September, October and November) in which prevalence was recorded (n=260/272) (95.5%), subsequent winter Dry season (December, January and February) with prevalence of (n=79/88) (89.7%) and then the relatively low prevalence was seen in summer (n=174/229) (75.98%). The result indicates that there is significance difference related to the season of the study P<0.05 (0.000). High seroprevalance was observed in medium herd size 92.3% (n=155/168) subsequently large herd size 86% (n=276/321) and in the last part small herd size 83 % (n = 83 / 100). The result indicates that there is no significance difference related to the herd size of the study P>0.05 (0.053) as shown in table 1.

This analysis also coincides with previous studies Camels in the larger herd size have slightly higher prevalence (n=324/347) (93.4%) than the small herd sized 92.3% (n=205/222), **[12]**. But the difference between the herd’s categories was not statistically significant the current study have little variation in the prevalence. Sero-prevalence of MERS-CoV in relation to production status was highly significant. With the study high prevalence was seen in lactating camels (n=145/146) (99.3%) following dry camels (158/163) (96.9%) consequently Pregnant camels (n=62/66) (93.9%) at the last N/A (young and Juveniles) Sero positivity indicates (n=149/214) (69.6%). In general, the result denotes that there is significant difference in sero positivity ratio among different production status of camels. The result indicates that there is significance difference P<0.05 as indicated in table 1 by which the juvenile with lactating camels may shed the virus and by transmitting the virus develop Sero positivity for MERS CoV.

Despite high Sero-positivity of MERS CoV antibody, the virus couldn’t be detected in the current study. This has been due to the development of MERS CoV antibody by large number of camels **[10]**. However in previous studies at Afar area (Fekadu *et al.*, 2017) (n= 7/100 (7%) of samples had detected by RT-PCR technique which was an indicative for the existence of circulating virus where it can be an evident for high sero positivity. Higher virus RNA detection rate in young animals compared with older animals could be related to a lack of prior immunity as published in previous studies in Saudi Arabia. Young animals were naïve and more susceptible to virus infection **[10]**.

## CONCLUSION

From the current study, there was clear evidence for overall high Sero-positivity of MERS-CoV in the study sites of Amibara district which was 87% (n=514/589). Among the study sites (Andido=45.16%, Melka Sedi 21.56 % and Angellele 20.54 %. Within the risk factors Age, Production status and season have significant difference in multivariate analysis for the prevalence of MERS CoV antibody.

The correlations of different risk factors were assessed in this study. In doing so, almost all risk factors were highly associated and were an important determinant for the disease In this study despite high Sero prevalence of MERS CoV antibody, the viral RNA is not able to be detected by RT-PCR test both at NAHDIC and HKU referral laboratories as previous studies indicated. This result disagree with the past studies high MERS CoV RNA rate detected in Ethiopia up to 15.7% ;(C.I. 95%, 8.2-28.0) **[10]**. In another study MERS CoV RNA with 7% was detected in Ethiopia between October 2014 and May 2015 **[12]**.

The possible causes for not getting /detecting the Viral RNA in the study area would be due to the following factors and challenges:-

Lack of sufficient information in understanding the viral shedding period or incubation time of the disease, lack of observation for apparent form of clinical sign of MERS CoV on camels as to enable taking the swab sample at early time of the disease, difficulty in deep swab sample taking process due to far distance of posterior turbinate of elongated nasal cavity of camels whereby it is the virus replication site compared to application swab stick length.

### Based on the above conclusion the following recommendations are forwarded

- Further study on the disease should be conducted in the study area by considering all aspects of the disease including in identifying other risk factors which will have value in the control of the disease.
- Even-though that, priority is given for swab sampling from nasal cavity of camel due to nature of replication site of virus; milk, urine and feces might be appropriate samples to detect the virus. Hence, these samples should be included at sampling.
- Camel abattoirs /slaughter houses to be included in taking swab samples from slaughtered camels to get access to the deep of nasal turbinate in getting the virus.
- A study to be considered by repeated swab sampling or as longitudinal study and focus of sampling to be given to well-marked and known juvenile and young camels as they are considered that, most of them are not developing MERS CoV antibody. This intensifies a chance of getting active virus to understand the virus characteristics.
- Since MERS CoV is one of the recently recognized zoonotic disease & camels are the sources of the virus to humans’ public awareness about the disease should be created in camel rearing pastoralist area.

## MATERIALS AND METHOD

### Study localities

A cross sectional study was carried out in Amibara districts of Afar region, Ethiopia Map of study sites for Amibara district Amibara sites as illustrated in figure 1. The district is located at latitude: 9° 39' N. Longitude 40°19'E within Administrative Zone three of Afar region bordered to the south by Awash Fentale district, to the west by Awash River which separates it from Dulecha, on the northwest by the Zone five administrative, to the north by Gewane, to the east by the Somali Region, and to the south east by Oromia Region. Amibara district has an average altitude of 867 m.a.s.l. Within the district, three study sites (Angellele, Melka Sedi, and Andido) were selected based on camel population density and being not previously studied.

**Figure 1:**
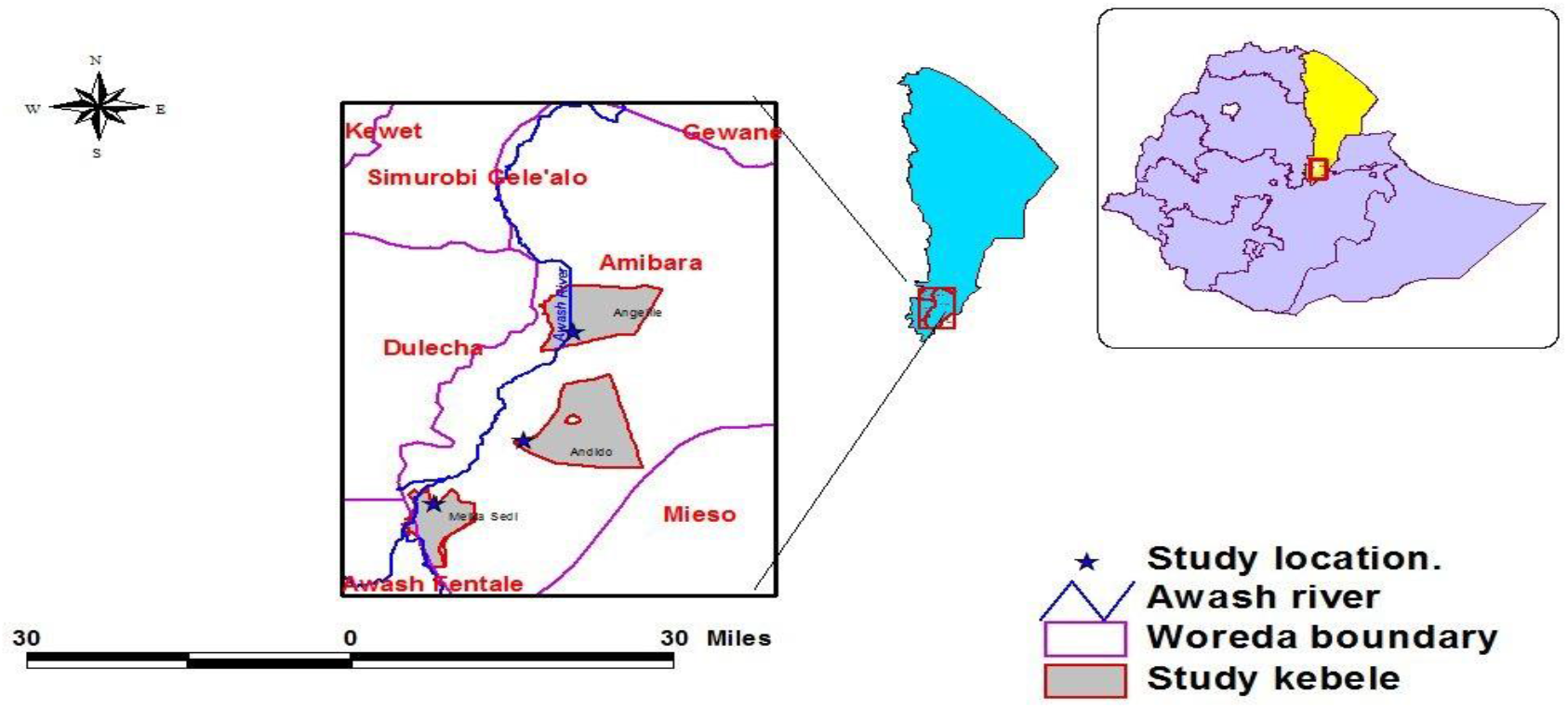
Map of study sites for Amibara district

### Study design and population

A cross sectional study design was used to assess the seroprevalance of MERS-CoV in Amibara. The target populations for the study were dromedary camel of all age groups, (juvenile, young and adult) and both sexes (male and female). Camel population in Amibara district was 148,769 **[14]**. The herd size of study population was composed of, high >30, medium =11-30 and low/small number of camel herds. =1-10 and the age categories is described as Juvenile <1 year, Young 1-3 years, Adult >3years **[15]**.

### Sample Size Determination and Sampling

The sample size determined for serological study was calculated by considering previously achieved epidemiological investigation of MERS-CoV with an expected prevalence of (92.3%) in the study area **[16]**. Thus, the calculated sample size using a 95% confidence interval at 5% absolute precision was 95% using the formula as described by **[17]**. The total sample size in camels was 110. Increasing the sample size was considered to increase the precision.

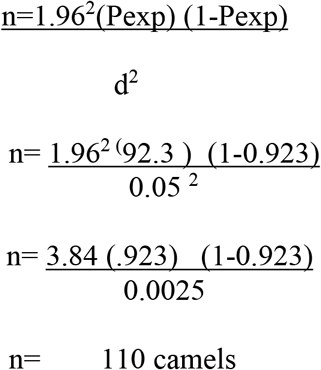

The increasing of sample size by 5 fold to enhance the precision of sampling and hence total sample was 589.

### Sampling techniques

Camels are restrained in all cases before sampling. Adequate health safety measures like wearing hand gloves, overall and mouth masks had been used at sampling site while sampling.

#### Blood sample for sera harvesting

Blood Samples were taken in duplicate from camels of each study three site. 10 ml of blood sample was collected from jugular vein using sterile needle and plain Vacutainer tube. The blood was allowed to clot at room temperature. Serum was separated from the clot by centrifugation at 3000rpm for 3 min and transferred to 2 ml cryo vial with a volume of 1.5-2 ml sera. The separated serum was labeled and kept under refrigeration (–20°C) until transported to NAHDIC for laboratory analysis both at NAHDIC and HKU. A total 589 sera were collected.

#### Nasal Swab sampling for detection of the virus

A total of 857 nasal swabs samples were collected in duplicate (for NAHDIC and HKU) by using applicator cotton swab **[18]**. The swab was taken for deep lateral turbinate. After taking sample, the swabs are immersed into 2 ml cryo vial containing 1.2 ml Viral transport medium (VTM) & preserved in liquid Nitrogen at −196 °C until transported to NAHDIC for keeping at −80°C freezer. Finally the swabs samples belonging to NAHDIC were tested in molecular laboratory and the other swab samples were shipped to HKU laboratory for MERS CoV RNA detection.

### Laboratory analysis

MERS CoV antibody detection through indirect ELISA test The MERS CoV antibody detection was carried out using the indirect ELISA test which is EUROIMMUN Anti −MERS-CoV S1 ELISA Camel (IgG) kit AG product of Lübeck, Germany according to manufacturer’s instructions **[19].**

#### Virus detection through RT-PCR

The Real-time polymerase chain reaction (RT-PCR) was used for detection of RNA of MERS-CoV. RNA extraction was carried out as described by the manufacturer instruction **[20].** Screening of the upstream of envelope gene (UpE) was done using UpE-FWD primer (GCAACGCGCGATTCAGTT) and UpE-Rev primer (GCCTCTACACGGGACCCATA) by reverse transcription quantitative PCR (RT-PCR) hydrolysis probe assay [**10]**.

### Data analysis

The Data obtained from the investigations was coded and stored in Excel spread sheets. The data was analyzed using STATA software version 15.0 software. Logistic regressions reporting the odd ratio at 95% confidence interval were used to determine the level of variation between the Sero-prevalence and the independent variable factors. The association of the explanatory and outcome variables was also analyzed by Chi^2^ test where p<0.05 indicates the significance level of the risk factors.

## ABREVATIONS

DNA: Deoxy ribonucleic acid
DPP: Dipeptidyl peptidase
E: Envelope protein
ELISA: Enzyme Linked ImmunoSorbent Assay
FAO: Food and Agriculture organization
HKU: Hong Kong University
IELISA: Indirect Enzyme Linked Immuno-Sorbent Assay
M: Matrix protein
MERS-CoV: Middle East Respiratory Syndrome Corona virus
N: Nucleocapsid protein
NAHDIC: National Animal Health Disease Investigation Centre
NSP: Non-structural protein
ORF: Open reading frame
P: Protein
RNA: Ribonucleic acid
RT-rtPCR: Reverse transcriptase real-time polymerase chain reaction
RT-PCR: Real time polymerase chain reaction
S: Spike-(surface glycoprotein)
SARS: Severe Acute Respiratory Syndrome
SP: Structural protein
UpE: Upstream Envelope
URT: Upper Respiratory Tract

## Declarations

### Ethics approval and consent to participate

Ethical clearance is approved and got permission from National Animal Health Diagnostic & Investigation Center (NAHDIC) Animal Research Scientific and Ethics Review Committee (ARSERC).

### Consent for publication

Not applicable

### Availability of data and materials

The data and materials are available.

## Competing interests

None of the authors of this paper have a financial or personal relationship with other people or organizations that could inappropriately influence or bias the content of this paper by any means.

## Funding

The study was sponsored by FAO, Ethiopia.

## Authors’ contributions

D.S contributed in sampling, epidemiological data gathering, laboratory tests, data acquisition, statistical analysis and drafting of the manuscript. F.A involved in the designing of the study, D.Z. Analysis of data and write up of the manuscript A.M. for lab test and analysis, G.M. contributed critical data analysis, interpretation and critical revision of the manuscript. E.W contributed participation in field work, data management and provision of logistics apart from these all authors included with in the manuscript contributed in editing and offering comment and approved the final version of the manuscript.

## Acknowledgements

The authors would like to thank National Animal Health and Disease Investigation Centre (NAHDIC) for allowing the laboratory facility in doing the tests, Food and Agricultural Organization (FAO), Ethiopia for the financial support for running costs of field activity and provision of laboratory materials and covering the costs for shipment and diagnostic costs of samples at H.K.U. Thanks also to H.K.U for generous cooperation of sample testing, analysis and offer a training in PCR and serology diagnostic techniques. Many thanks to Amibara Agricultural and Livestock staffs Dr Mekonnen, Seid, Million and Hummed for their cooperation in mobilizing and convincing camel owners of the pastoral community. In sample collection from camels.

## Notes

### Competing Interest Statement

The authors have declared no competing interest.

